# Association between Neighborhood Socioeconomic Status and Executive System Activation in Youth

**DOI:** 10.1101/2021.08.12.454903

**Authors:** Kristin Murtha, Bart Larsen, Adam Pines, Linden Parkes, Tyler M. Moore, Azeez Adebimpe, Aaron Alexander-Bloch, Monica E. Calkins, Diego G. Davila, Martin A. Lindquist, Allyson P. Mackey, David R. Roalf, J. Cobb Scott, Daniel H. Wolf, Ruben C. Gur, Raquel E. Gur, Ran Barzilay, Theodore D. Satterthwaite

## Abstract

Low socioeconomic status has been shown to have detrimental effects on cognitive performance, including working memory (WM). As executive systems that support WM undergo functional neurodevelopment during adolescence, environmental stressors at both individual and community levels may have a particularly strong impact on cognitive outcomes. Here, we sought to examine how neighborhood socioeconomic status (SES) impacts task-related activation of the executive system during adolescence, and to determine whether this effect mediates the relationship between neighborhood SES and WM performance. To address these questions, we studied 1,158 youths (age 8-22) that completed a fractal *n*-back WM task during fMRI at 3T as part of the Philadelphia Neurodevelopmental Cohort. We found that higher neighborhood SES was associated with greater activation of the executive system to WM load, including the bilateral dorsolateral prefrontal cortex, posterior parietal cortex, and precuneus. These associations remained significant when controlling for related factors like parental education and exposure to traumatic events. Furthermore, high dimensional multivariate mediation analysis identified two distinct patterns of brain activity within the executive system that significantly mediated the relationship between neighborhood SES and task performance. Together, these findings underscore the importance of the neighborhood environment in shaping executive system function and WM in youth.

---

Low socioeconomic status (SES) has detrimental effects on diverse measures of brain development and cognition in youth (Bradley and Corwyn 2002; Pollak and Wolfe 2020). In particular, working memory (WM) has been consistently shown to vary by SES, with low-SES individuals having poorer performance (Noble et al. 2007; Hackman et al. 2015; Leonard et al. 2015). The impact of SES on WM may be particularly important during youth, when WM improves dramatically (Gur et al. 2012; Satterthwaite et al. 2013; Gur et al. 2014; Ullman et al. 2014; Luna et al. 2015; Simmonds et al. 2017). Indeed, prior work suggests that differences in WM may in part explain SES-related differences in academic performance (Gathercole et al. 2004; Best et al. 2011). However, the neurobiological mechanisms that link low SES to WM deficits in youth remain incompletely described (Hart et al., 2007; Rosen et al., 2019; Sheridan & McLaughlin, 2014).

Studies have provided evidence that WM performance is subserved by a spatially distributed set of brain regions involved in executive function, including the dorsolateral prefrontal cortex (DLPFC), anterior insula, intraparietal sulcus, precuneus, and cerebellum (Satterthwaite et al. 2013; Samartsidis et al. 2019; Rosenberg et al. 2020). During WM tasks, activation of this distributed executive system has been shown to be reduced in youth with low family income (Finn et al. 2017), while lower parental education has also been associated with inefficient recruitment of elements of the executive system (Sheridan et al. 2017). Additionally, higher levels of early life stress, including measures of financial strain and housing challenges, predict greater resting-state homogeneity in the left middle frontal gyrus, which in turn is associated with lower levels of cognitive control (Demir-Lira et al. 2016). These results from fMRI cohere with findings from studies using structural MRI, which have reported that low-SES youth have reduced cortical surface area in executive regions (Noble et al. 2015; Leonard et al. 2019).

Notably, most studies examining the neural underpinnings of the relationship between SES and cognitive function have operationalized SES using individual-specific variables, such as family income or parental education. However, accruing evidence suggests that neighborhood-level factors can impact cognition above and beyond individual-specific measures (Brito and Noble 2014; Tomlinson et al. 2020). Specifically, recent work has begun to utilize measures of environmental adversity at the neighborhood level, which include crime rates, social capital, or access to housing (Leventhal and Brooks-Gunn 2000). These factors have been shown to impact both physical (Boylan and Robert 2017) and mental health outcomes (Aneshensel and Sucoff 1996).

Work using large-scale neuroimaging studies has found environmental adversity measured on the neighborhood scale is associated with neurocognitive performance across a variety of domains, including executive function and WM (Gur et al. 2019; Vargas et al. 2020). Neighborhood disadvantage has also been related to other neuroimaging parameters, including lower gray matter volume (Butler et al. 2018), BOLD activation in response to social exclusion (Gonzalez et al. 2015), development of functional networks (Tooley et al. 2019), as well as structural differences including lower surface area in the prefrontal cortex (Vargas et al. 2020) and lower volume in the dorsolateral prefrontal cortex and right hippocampus (Taylor et al. 2020). These studies suggest that understanding the effects of SES on cognition and brain function requires consideration of adversity measured at the community level. However, no prior studies have examined the relationship between neighborhood SES and executive system activation during WM.

In the current study, we investigated how neighborhood SES impacts WM and brain function during youth. Specifically, we used geocoded block-level data to assess neighborhood-level SES and investigate the association between SES and executive system activation during a WM fMRI task. We hypothesized that lower neighborhood SES would be associated with reduced activation of the executive system, and that multivariate patterns of brain activation within these regions would mediate the relationship between neighborhood SES and performance on an in-scanner WM task.

## MATERIALS AND METHODS

### Participants

We examined a cross-sectional sample of 1,536 participants from the Philadelphia Neurodevelopmental Cohort (PNC) (Satterthwaite et al. 2016) who underwent functional neuroimaging while completing a fractal *n*-back task (mean age=14.9, range 8-23, 837=female). Of these individuals, 378 were excluded for medical comorbidities that impact brain function (n=148), image quality (n=227), or incomplete clinical data (n=3). The final sample included in the analysis consisted of 1,158 individuals (mean age=15.4, range 8-23; 622=female).

### Measure of Neighborhood Socioeconomic Status

The quantification of neighborhood socioeconomic status in this sample has been detailed previously (Moore et al. 2016). A set of geocoded variables were obtained from participant addresses and incorporated 2010 census data from the greater Philadelphia area. Examples of characteristics in this census-block level data included median family income, percent of residents who are married, percent of homes which are family homes, and percent of people in poverty. Census block groups typically contain 600-3,000 persons and can also vary in square footage, meaning they can vary (sometimes extremely) in density. A weighted factor score of neighborhood level SES was generated from these variables using the Thurstone Method (Thurstone 1935).

### Task Design

The fractal *n-*back task used in the PNC has previously been described in detail (Satterthwaite et al. 2012; Shanmugan et al. 2016). Briefly, participants completed a fractal *n*-back task (Ragland et al. 2002) during fMRI as a measure of WM (**Figure 1A**). The task was structured with a block-design using 3 conditions of increasing WM load: 0-back, 1-back, and 2-back. In the 0-back condition, participants responded to a single target image. In the 1-back condition, participants responded if the image presented was the same as the previous image. In the 2-back condition, participants responded if the image presented was the same as the image presented 2 trials previously. Each condition included 20 trials over 60s, and was repeated over 3 blocks. Participants were cued with verbal instructions as to which condition they would be completing at the beginning of each block.

**Figure 1:**
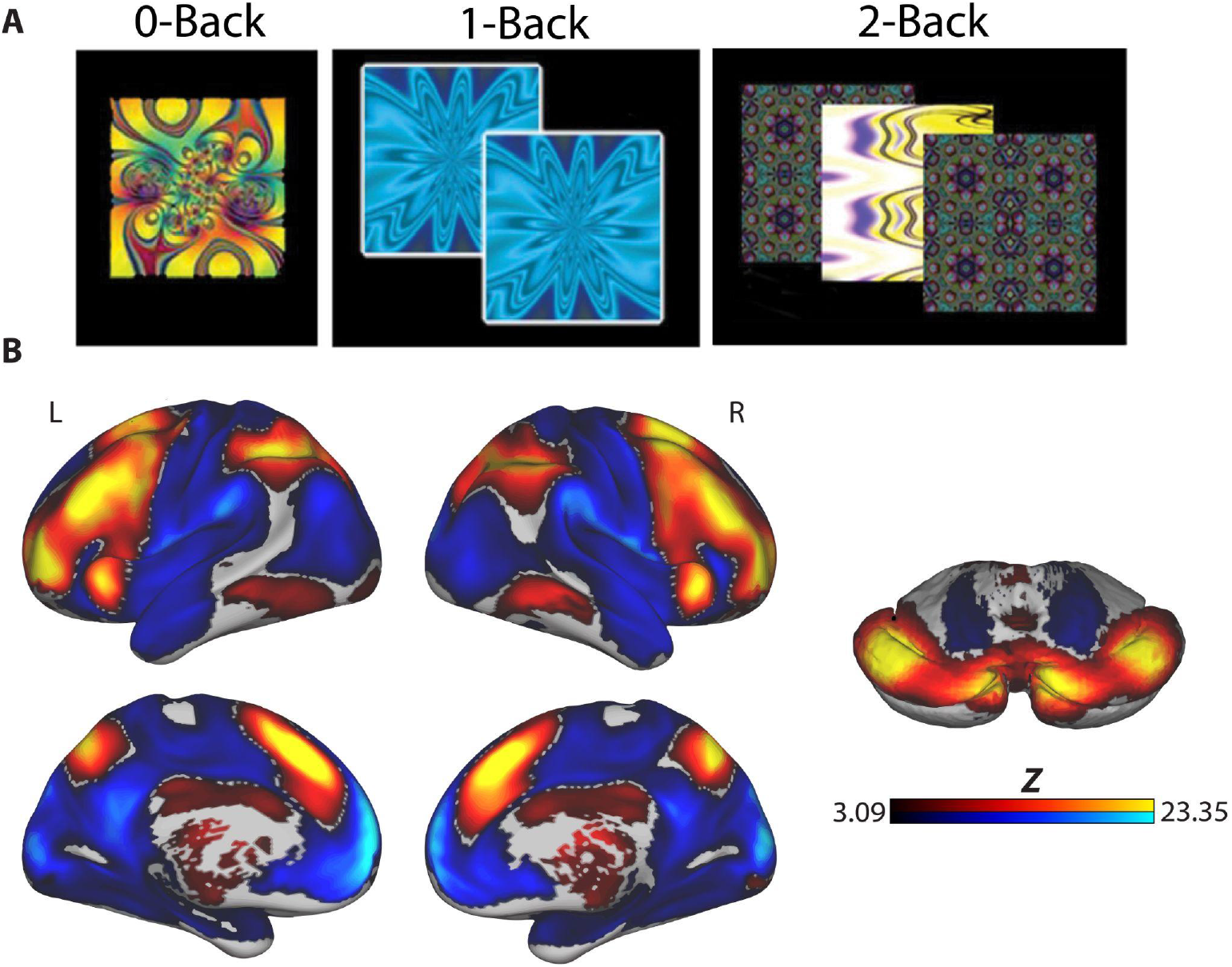
The effect of working memory load on brain activation. **(A)** Executive function was measured using a fractal version of the *n*-back WM task. **(B)** Increased working memory load, operationalized by the 2-back>0-back contrast, robustly activated the distributed executive system, while deactivating non-executive regions (image was thresholded at *z*>3.09, cluster corrected at *p*<0.05).

### Image Acquisition

Participants completed a neuroimaging protocol that included fMRI, T1 and B0 sequences, collected at a single scanner (Siemens 3-T, 32-channel head coil). A magnetization-prepared rapid acquisition gradient echo T1-weighted image was acquired to aid spatial normalization to standard atlas space, using the following parameters: TR, 1810 ms; TE, 3.51 ms; TI, 1100 ms; FOV, 180 × 240 mm; matrix, 192 × 256; 160 slices; slice thickness/gap, 1/0 mm; flip angle, 9°; effective voxel resolution, 0.9 × 0.9 × 1 mm. Blood oxygen level-dependent fMRI was acquired using a whole-brain, single-shot, multislice, gradient-echo echo-planar sequence with the following parameters: 231 volumes; TR, 3000; TE, 32 ms; flip angle, 90°; FOV, 192 × 192 mm; matrix 64 × 64; 46 slices; slice thickness/gap 3/0 mm; effective voxel resolution, 3.0 × 3.0 × 3.0 mm. Additionally, a B0 field map was acquired for application of distortion correction procedures, using the following double-echo gradient recall echo sequence: TR, 1000 ms; TE1, 2.69 ms; TE2, 5.27 ms; 44 slices; slice thickness/gap, 4/0 mm; FOV, 240 mm; effective voxel resolution, 3.8 × 3.8 × 4 mm.

### Image Processing

Image preprocessing is described in detail in our prior work describing this dataset (Satterthwaite et al. 2014; Shanmugan et al. 2016). Briefly, basic pre-processing of *n*-back task images used tools from FSL, including slice time correction, skull stripping, motion correction, spatial smoothing, and grand mean scaling. Time-series analysis of subject-level imaging data modeled the *n*-back task’s three condition blocks (0-back, 1-back, and 2-back) using FEAT. Subject-level statistical maps of the primary 2-back>0-back contrast were distortion corrected, coregistered to the MNI 152 1-mm template registered T1 image, and normalized using Advanced Normalization Tools (Avants et al. 2008). Images were downsampled to 2 mm resolution before group-level analysis. All transforms were concatenated so that only a single interpolation was performed.

### Behavioral Analysis

We summarized performance during the *n*-back task using the signal detection measure *d’*. This measure incorporates both correct responses and false positives in order to limit the impact of response bias on the measure of accuracy (Snodgrass and Corwin 1988).

### Group-Level Analysis

Our primary group-level analysis sought to characterize the association between neighborhood SES and changes in brain activation under WM load (2-back>0-back). We conducted a mass-univariate voxelwise analyses using tools from FSL (Jenkinson et al. 2012), where activation in the 2-back vs. 0-back condition was the outcome, and neighborhood SES was the predictor of interest; age, sex, and in-scanner motion were included as covariates. We controlled for multiple comparisons using cluster correction as implemented in FSL (voxel height z > 3.09, cluster probability *p* <0.05). Visualizations were generated using Connectome Workbench, developed under the auspices of the Human Connectome Project at Washington University in St. Louis and associated consortium institutions (http://www.humanconnectome.org) (Marcus et al. 2011).

### Sensitivity Analyses

To evaluate the potentially confounding influence of other participant factors, we conducted sensitivity analyses that included additional model covariates. Specifically, we repeated the mass-univariate analysis described above, but also included parental educational attainment, exposure to traumatic stress, and task performance (*d’*) as model covariates. Traumatic stress was assessed as part of a structured clinical interview (GOASSESS), and quantified by the lifetime number of categories of exposure to traumatic stressful events experienced by a participant (range 0-8) (Calkins et al. 2015; Barzilay et al. 2019).

### Mediation Analyses

As a final step, we sought to understand how multivariate patterns of brain activation might mediate the observed association between neighborhood SES and WM performance. To do this, we examined principal directions of mediation (Chén et al. 2018; Geuter et al. 2020) using the M3 Mediation toolbox from the Cognitive and Affective Neuroscience Lab (CANlab; available at https://github.com/canlab/MediationToolbox). This type of mediation analysis seeks to extract linear combinations of high-dimensional neuroimaging data that maximize the indirect effect (i.e., the mediation) between independent and dependent variables. The result is a set of orthogonal principal directions of mediation (PDMs) that can be mapped back to the original neuroimaging data space to provide interpretable mediation effects (see Chén et al. for details). To avoid overfitting and test the generalizability of the PDMs observed in our data, we first divided our sample into training (*n* = 580) and testing (*n* = 578) sets that were matched on neighborhood SES. This enabled us to model PDMs in the training set and then apply the model to the unseen data in our test set.

Prior to estimating PDMs, we used singular value decomposition to reduce the dimensionality of the neuroimaging data from the training subset. We selected all principal components that explained greater than 1% of variance in the data, which yielded a total of 10 components. The PDM’s were then estimated using this 580 (participants) x 10 (principal components) reduced matrix. Next, significance of the indirect path associated with each PDM was determined using a bootstrap procedure with 10,000,000 iterations, controlling for age, sex and motion as nuisance covariates. To validate the estimated PDM’s, we applied the model generated on the training data to the unseen test data and performed the same bootstrapping procedure to assess significance of indirect path coefficients while controlling for the same set of nuisance covariates. Thus, the mediation effect encoded by each PDM was tested for significance twice; once in the training data and again on unseen test data. Next, in order to interpret the mediation effects, we extracted the voxel-weights from the PDM’s that yielded significant indirect paths on both the training and the testing data. Significance of each PDM was assessed using bootstrap analyses to derive significance at voxel level, while controlling for type I error with cluster correction as stated above (voxel height z > 3.09, cluster probability *p* < 0.05).

## RESULTS

### Lower neighborhood socioeconomic status is associated with attenuated activation of the executive system

As previously reported (Satterthwaite et al. 2013), the *n*-back task robustly activated the distributed components of the brain’s executive system (**Figure 1B**) and deactivated non-executive regions, including the default mode network. We hypothesized that higher neighborhood socioeconomic status (SES) would be associated with greater recruitment of the executive system. Mass univariate voxel-wise analysis revealed that higher neighborhood SES was associated with greater bold activation in three large clusters within the executive system (see **Figure 2** and **Table 1**). The first cluster included bilateral dorsolateral prefrontal cortex (DLPFC), anterior insula, paracingulate, frontal pole, and the supplementary motor area. The second cluster spanned parietal cortex and cerebellum, including bilateral superior parietal cortex, precuneus, and bilateral cerebellar crus I & II. The third cluster included parts of the inferior temporal cortex and temporal pole. Given the large size of each cluster, we increased the statistical threshold (z>4.0, cluster probability *p*<0.05) in order to break up large clusters and increase the specificity of the reported cluster coordinates. This process yielded 13 clusters that are presented in **Table 2**.

**Figure 2:**
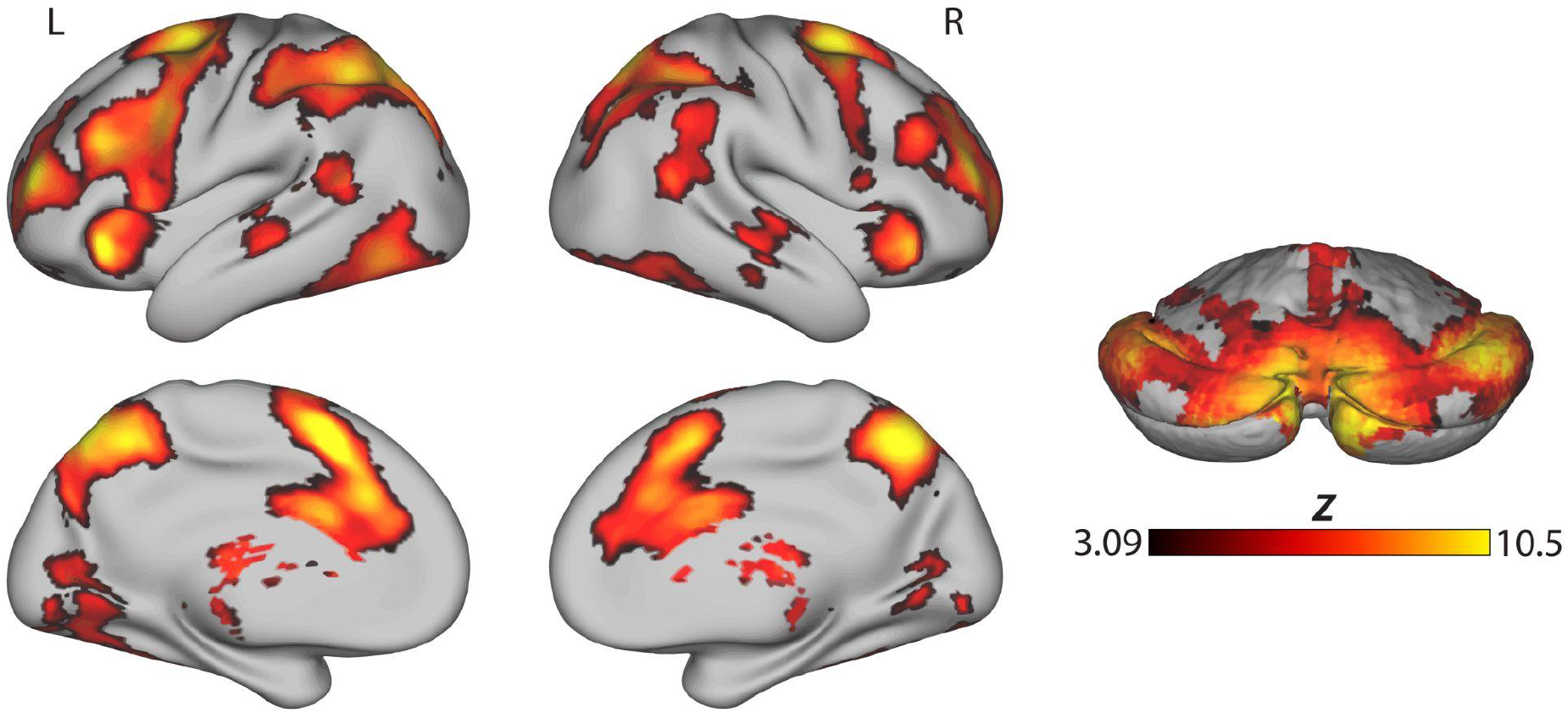
Higher neighborhood SES was significantly associated with greater activation throughout the executive system. A mass univariate analysis revealed 3 large clusters of significant activation (voxel height *z*>3.09, cluster probability *p*<0.05). The first was a large prefrontal cluster that included bilateral dorsolateral prefrontal cortex, anterior insula, paracingulate, and frontal pole, and the bilateral thalamus. The second was a large parietal cluster that included bilateral superior parietal cortex and precuneus, bilateral temporoparietal junction, right posterior temporal cortex, as well as bilateral cerebellar crus I & II. Finally, the third cluster included the temporal cortex and temporal pole. Cluster coordinates are listed in Tables 1 and 2.

**Table 1.** Association of SES and *n*-back on the 2-back vs. 0-back conditions (a priori statistical threshold)*

| Cluster Region | Voxels | Cluster Probability | Max (Z) | Peak X (mm) | Peak Y (mm) | Peak Z (mm) |
| --- | --- | --- | --- | --- | --- | --- |
| Large bilateral prefrontal cluster | 20817 | <0.0001 | 10.5 | -30 | -4 | 66 |
| Large bilateral parietal and cerebellar cluster | 17870 | <0.0001 | 10.5 | -12 | -76 | 58 |
| Cluster in temporal cortex and temporal pole | 485 | 0.0401 | 4.53 | 44 | -30 | -10 |

\*minimum voxel height $z > 3.09$ , cluster probability $p < 0.05$ , reported LPI.

**Table 2.** Association of SES and *n*-back on the 2-back vs. 0-back conditions (elevated statistical threshold)*

| Cluster Region | Voxels | Cluster Probability | Max (Z) | Peak X (mm) | Peak Y (mm) | Peak Z (mm) |
| --- | --- | --- | --- | --- | --- | --- |
| Large prefrontal cluster, including bilateral dorsolateral prefrontal cortex, anterior insula, paracingulate, and frontal pole | 12043 | <0.0001 | 10.5 | -30 | -4 | 66 |
| Large parietal cluster, including bilateral superior parietal cortex and precuneus | 7432 | <0.0001 | 10.5 | -12 | -76 | 58 |
| Large cerebellar cluster, including bilateral crus II and left crus I | 1581 | <0.0001 | 8.38 | -4 | -78 | -30 |
| Large cluster including right cerebellar crus I and right posterior temporal cortex | 1130 | <0.0001 | 7.48 | -36 | -40 | -38 |
| Bilateral thalamus | 654 | <0.0001 | 6.2 | 8 | -24 | 14 |
| Lingual gyrus | 217 | 0.0026 | 5.62 | -12 | -64 | 0 |
| Left temporoparietal junction | 91 | 0.0142 | 5.41 | -52 | -54 | 16 |
| Occipital fusiform gyrus | 79 | 0.0172 | 4.83 | 40 | -62 | -12 |
| Right temporoparietal junction | 76 | 0.0181 | 5.25 | 58 | -48 | 16 |
| Lateral temporal cortex | 61 | 0.0235 | 4.53 | 46 | -26 | -2 |
| Lateral temporal cortex | 53 | 0.0273 | 4.54 | -46 | -30 | -2 |
| Temporal fusiform cortex | 39 | 0.0360 | 4.91 | -34 | -44 | -20 |
| Intracalcarine cortex | 28 | 0.0459 | 4.64 | 16 | -68 | 8 |

\* voxel height $z > 4.0$ , cluster probability $p < 0.05$

Next, we conducted post-hoc analyses to understand the contribution of WM during the task to the observed associations with SES. Analysis of activation during the 0-back and 2-back conditions contrasted with baseline suggested that the increased activation in the executive system as a result of neighborhood SES was driven primarily by the 2-back condition. Separate linear models conducted in the first cluster, including prefrontal regions, and second cluster, including parietal regions, revealed significant relationships between neighborhood SES and 2-back activation, whereas neighborhood SES was not significantly associated with activation in the 0-back condition (**Figure 3**).

**Figure 3:**
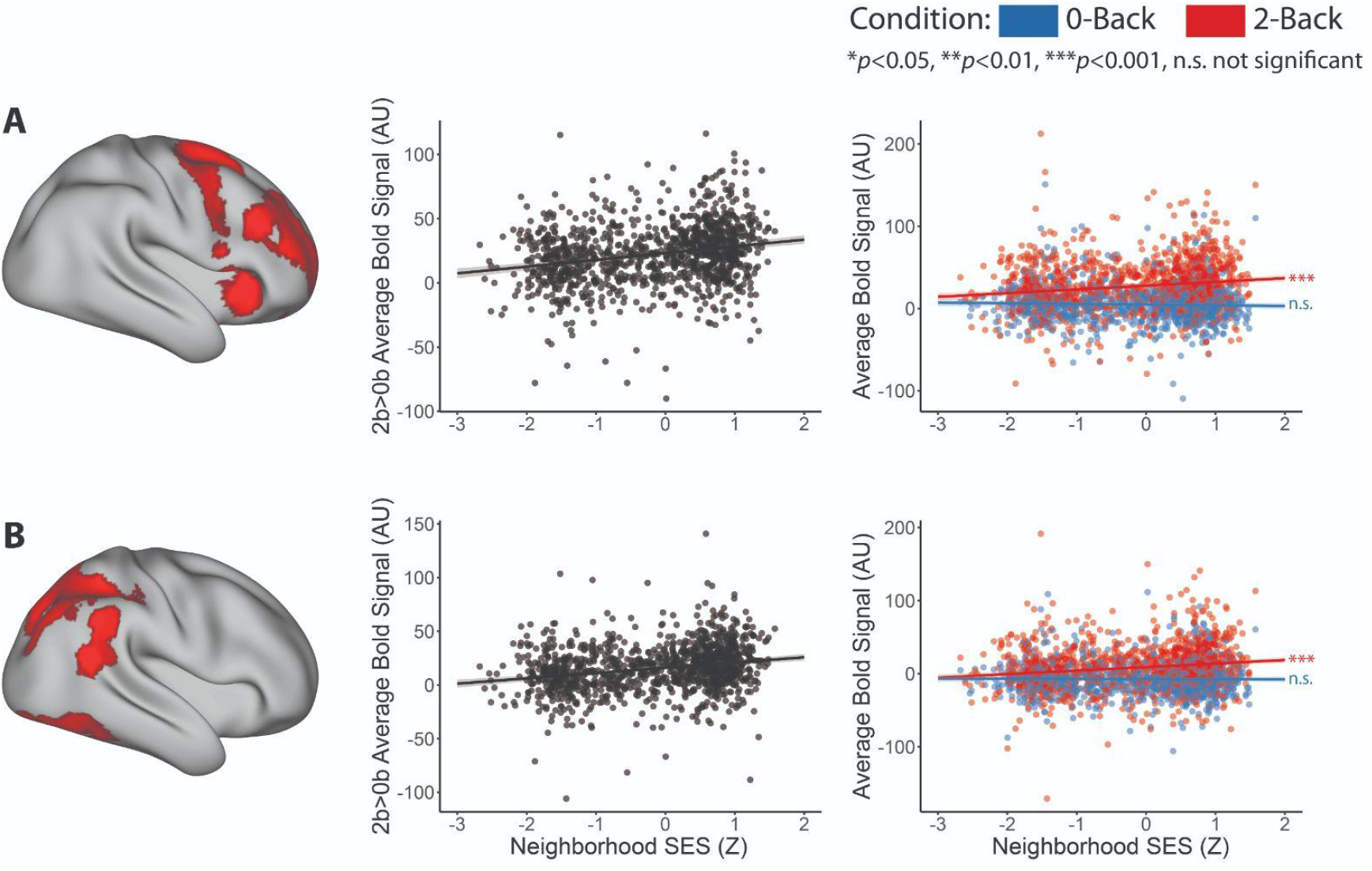
The effect of neighborhood SES on executive system activation was driven by individual differences in activation during the high working memory load condition (2-back). In both the prefrontal cluster **(A)** and parietal cluster **(B)**, post-hoc linear models of the effect of neighborhood SES on activation during the 2-back and 0-back conditions each revealed significant relationships between SES and 2-back activation *(p<*0.0001). In contrast, neighborhood SES was not significantly associated with activation during the 0-back condition.

### Sensitivity analyses

Sensitivity analyses revealed that the association between neighborhood SES and activation during the *n*-back task remained significant when controlling for additional covariates. A separate mass univariate voxelwise analysis that controlled for additional covariates including in-scanner task performance (as measured by *d’*), exposure to traumatic events, as well as paternal and maternal education, revealed a highly convergent pattern of results (**Table 3**). These findings suggest that neighborhood SES is associated with individual differences in executive activation over and above other commonly used measures of life experience and is not due to individual differences in task performance during the scan.

**Table 3.** Sensitivity Analysis including maternal and paternal education, traumatic stress exposure, and task performance as additional covariates (n=1059)*

| Cluster Region | Voxels | Cluster Probability | Max (Z) | Peak X (mm) | Peak Y (mm) | Peak Z (mm) |
| --- | --- | --- | --- | --- | --- | --- |
| Large parietal cluster, including bilateral superior parietal cortex and precuneus | 1568 | <0.0001 | 5.99 | 8 | -66 | 64 |
| Left prefrontal cluster | 977 | <0.0001 | 5.16 | -30 | -4 | 66 |
| Right prefrontal cluster | 694 | 0.0003 | 4.84 | 30 | 2 | 68 |
| Left dorsolateral prefrontal cortex | 485 | 0.0021 | 4.89 | -30 | 46 | 42 |

\* voxel height $z > 3.09$ , cluster probability $p < 0.05$

### Multivariate patterns of activation mediate the relationship between neighborhood socioeconomic status and working memory performance

Given the observed association between neighborhood SES and executive activation, the significant association between neighborhood SES and task performance (**Figure 4**), and the known relationship between neural activation and WM performance (Shamosh et al. 2008; Satterthwaite et al. 2013), we investigated whether multivariate patterns of brain activation mediated the relationship between neighborhood SES and task performance. We first conducted the mediation analysis in a training sample of 580 participants (**Figure 5A)**. For each PDM, we estimated *a, b* and *ab* path coefficients representing the relationship between neighborhood SES and brain activation, brain activation and task performance, and the mediation effects, respectively. The absolute *ab* path coefficients represent the extent to which activation mediates the relationship between neighborhood SES and task performance. In our training sample, bootstrap analysis revealed that 5 PDM’s had significant *ab* paths after FDR correction (*p-train*_*fdr*_<0.05), and their combined *ab* path coefficients explained 15.34% of the variance of the relationship between neighborhood SES and task performance.

**Figure 4:**
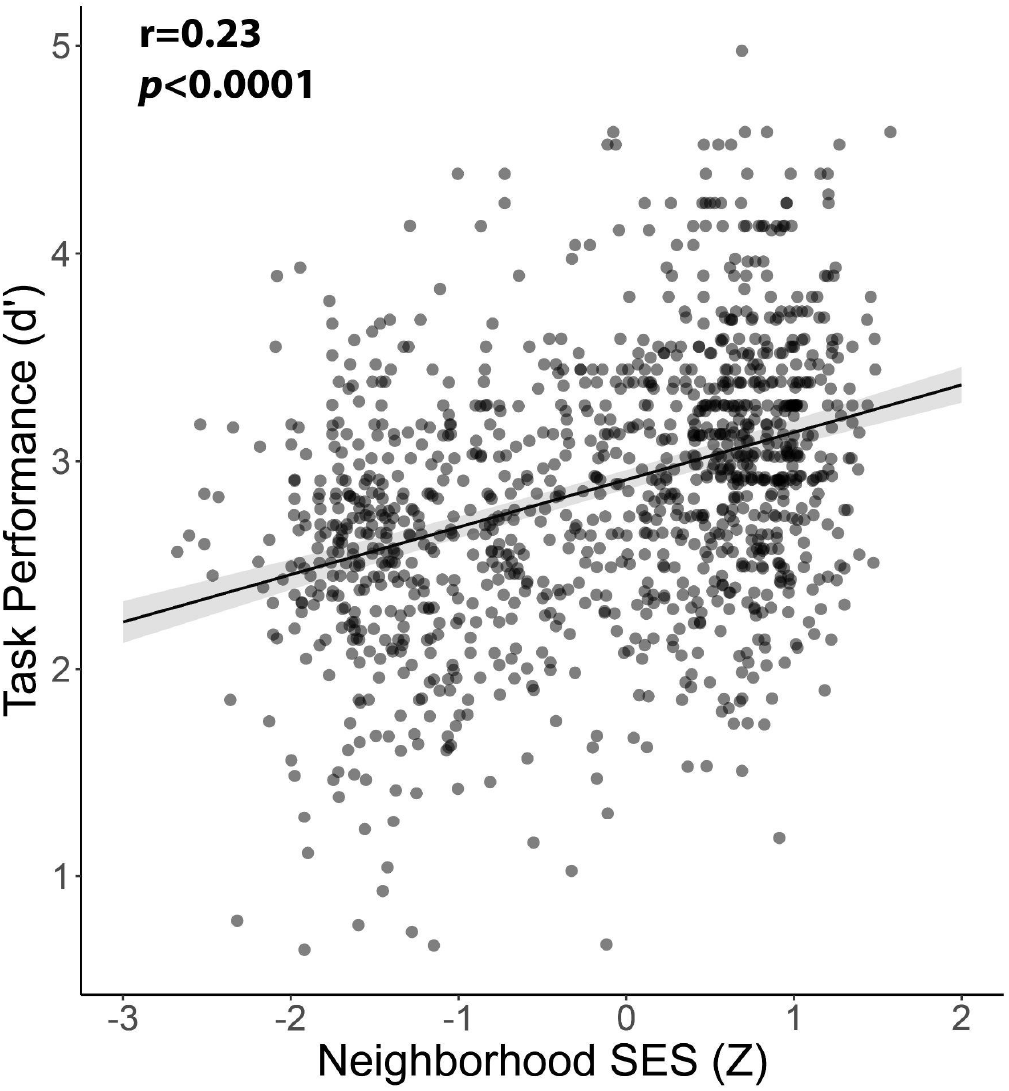
Neighborhood SES is positively associated with WM task performance. WM performance was quantified as *d’* across all *n*-back conditions, while covarying for age and sex.

**Figure 5:**
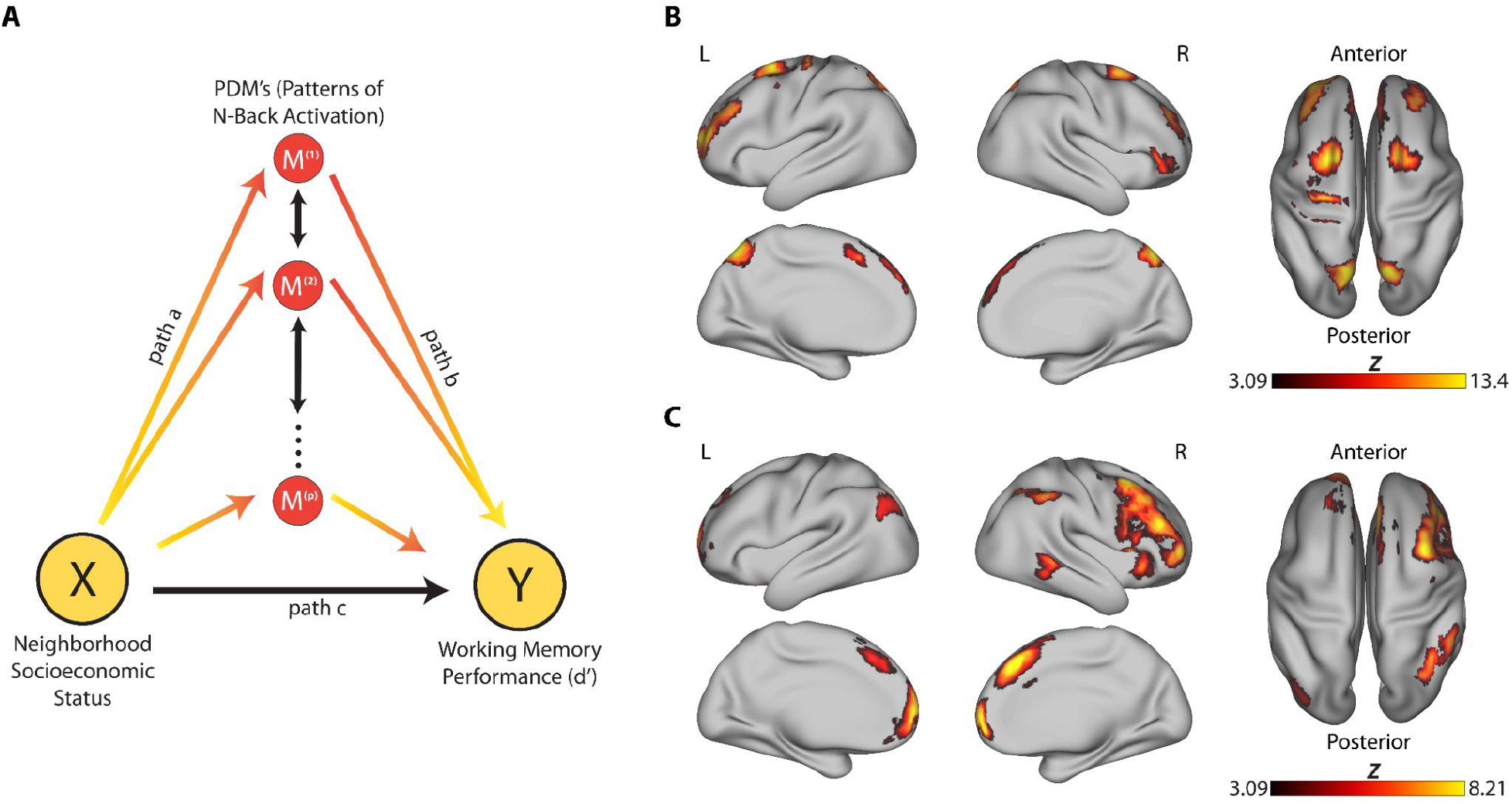
Multivariate patterns of executive system activation mediate the relationship between neighborhood SES and working memory performance. **(A)** We assessed whether multivariate patterns of activation mediated the relationship between neighborhood SES and working memory performance using high-dimensional mediation analysis. Yellow circles represent independent (neighborhood SES) and dependent variables (working memory performance). Red circles represent potential Principal Directions of Mediation (PDM’s), or patterns of brain activation that mediate the relationship between the dependent and independent variables. This analysis revealed 2 significant patterns of brain activation that mediated the relationship between neighborhood SES and task performance (*p-train*_*fdr*_<0.01, *p-test*_*fdr*_<0.001). **(B)** PDM1 included regions in the medial prefrontal cortex, left and right DLPFC, as well as the middle frontal and superior frontal and precentral gyri. **(C)** PDM2 included the superior frontal gyrus, right middle frontal gyrus and frontal pole, as well as the left frontal pole, left lateral occipital cortex, and middle temporal gyrus. Multiple comparisons were accounted for using cluster correction with a voxel height of *z*>3.09 and cluster probability *p*<0.05.

Next, we applied the PDM’s generated in our training data to a held-out test set of 578 participants and reperformed bootstrap analysis of the absolute *ab* paths. The *ab* paths of the first 2 PDM’s remained significant in the left out sample and survived FDR correction (*p-test*_*fdr*_<0.05; **Table 4**) while controlling for age, sex, and in-scanner motion. These results suggest that the patterns of brain activation encoded in PDMs 1 and 2 not only partially mediate the relationship between neighborhood SES and WM, they also robustly generalize to unseen data. Both the *a* and *b* path coefficients were positive (note that *a* paths are always fixed as positive by this method), meaning that higher SES was linked to more activity in the brain regions associated with each PDM, and that activation in these regions was also positively correlated to better WM performance.

**Table 4.** Path Coefficients, Z-scores, and bootstrapped p-values for train and test data in PDM’s 1 & 2.

|  | Train |  |  | Test |  |  |
| --- | --- | --- | --- | --- | --- | --- |
| | coeff | $z$ | $p$ | coeff | $z$ | $p$ |
| PDM 1 |  |  |  |  |  |  |
| a | 924.39011 | 5.22 | <0.0001 | 1075.65 | 5.16 | <0.0001 |
| b | 0.00010 | 5.04 | <0.0001 | 0.00 | 5.21 | <0.0001 |
| c' | 0.09160 | 4.34 | <0.0001 | 0.17 | 5.24 | <0.0001 |
| c | 0.18421 | 5.23 | <0.0001 | 0.27 | 5.28 | <0.0001 |
| ab | 0.09261 | 5.32 | <0.0001 | 0.10 | 5.26 | <0.0001 |
| PDM 2 |  |  |  |  |  |  |
| a | 438.74657 | 3.51 | 0.0004 | 608.48 | 5.22 | <0.0001 |
| b | 0.00007 | 5.18 | <0.0001 | 0.00 | 5.08 | <0.0001 |
| c' | 0.15340 | 5.23 | <0.0001 | 0.23 | 5.21 | <0.0001 |
| c | 0.18420 | 5.23 | <0.0001 | 0.27 | 5.28 | <0.0001 |
| ab | 0.03080 | 3.6 | 0.0003 | 0.04 | 5.36 | <0.0001 |

As a final step, we examined the spatial distribution of the significant PDMs (**Table 5**). The first PDM included clusters in the medial prefrontal cortex, left and right DLPFC, as well as the middle frontal and superior frontal and precentral gyri (**Figure 5B**). The second PDM included a large right-predominant cluster including the superior frontal gyrus, right middle frontal gyrus and frontal pole, as well as clusters in the left frontal pole, left lateral occipital cortex, and middle temporal gyrus (**Figure 5C**). These findings suggest that the association between neighborhood SES and WM performance may in part be mediated by differences in executive system activation.

**Table 5.** Mediation Results*.

| PDM1 |  |  |  |  |  |  |
| --- | --- | --- | --- | --- | --- | --- |
| Cluster Region | Voxels | Cluster Probability | Max (Z) | Peak X (mm) | Peak Y (mm) | Peak Z (mm) |
| Medial prefrontal cortex | 1916 | <0.0001 | 8.21 | 6 | 60 | 44 |
| Precuneus, lateral occipital cortex | 1585 | <0.0001 | 13.4 | -16 | -74 | 56 |
| Left dorsolateral prefrontal cortex | 1517 | <0.0001 | 8.21 | -30 | 64 | 16 |
| Right dorsolateral prefrontal cortex | 780 | <0.0001 | 7.94 | 34 | 46 | 40 |
| Premotor & parietal cortex, middle frontal gyrus | 508 | <0.0001 | 8.21 | -28 | 4 | 66 |
| Dorsal premotor parietal cortex, superior frontal Gyrus | 384 | <0.0001 | 6.82 | 28 | 4 | 64 |
| Primary motor cortex, precentral gyrus | 361 | <0.0001 | 7.14 | -30 | -24 | 72 |
| Right frontal pole | 287 | 0.0003 | 5.13 | 48 | 50 | -4 |
| Paracingulate gyrus | 179 | 0.0055 | 5.37 | -2 | 16 | 48 |
| PDM2 |  |  |  |  |  |  |
| Cluster Region | Voxels | Cluster Probability | Max (Z) | Peak X (mm) | Peak Y (mm) | Peak Z (mm) |
| Right dorsolateral prefrontal cortex | 5601 | <0.0001 | 8.21 | 2 | 40 | 54 |
| Ventromedial prefrontal cortex | 2332 | <0.0001 | 8.21 | -4 | 64 | -2 |
| Superior parietal cortex | 1074 | <0.0001 | 8.21 | 44 | -52 | 58 |
| Middle frontal gyrus | 399 | <0.0001 | 7.59 | -54 | 14 | 34 |
| Lateral occipital cortex | 376 | 0.0001 | 6.27 | -46 | -74 | 38 |
| Left frontal pole | 181 | 0.0099 | 6.38 | -42 | 54 | 2 |
| Superior temporal sulcus | 172 | 0.0125 | 3.95 | 66 | -38 | 2 |

\* voxel height $z > 3.09$ , cluster probability $p < 0.05$

## DISCUSSION

We found that higher neighborhood SES is associated with greater activation of the executive system, including the bilateral prefrontal cortex and superior parietal cortex. Our findings indicate that SES measured at the neighborhood level is related to executive system recruitment over and above other related factors, such as parental education level and exposure to traumatic events. Furthermore, we demonstrate that multivariate patterns of executive system activation partially mediate the relationship between neighborhood-level SES and performance on a WM task. These results provide novel evidence that neighborhood characteristics may influence WM performance through their impact on the brain’s executive system.

Higher neighborhood SES was associated with increased activation in brain regions associated with WM performance (Crone et al. 2006; Satterthwaite et al. 2013; Samartsidis et al. 2019; Rosenberg et al. 2020). Specifically, SES was associated with activation across regions including the bilateral DLPFC, paracingulate cortex, bilateral superior parietal cortex, and precuneus. These findings expand on earlier work in the same sample, showing reduced regional homogeneity and amplitude of low-frequency fluctuations during resting state fMRI in frontoparietal regions associated with neighborhood SES (Gur et al. 2019; Tooley et al. 2019). The current work also aligns with findings that higher family SES, measured by income-to-needs ratio, is associated with increased BOLD activation in the prefrontal cortex during WM tasks (Rosen et al. 2018).

Notably, we found that our observed pattern of effects was driven by the 2-back task condition, where the greatest WM load was present. This suggests that the relationship between SES and executive system activation is most evident in more cognitively demanding task contexts. No significant relationship between activation and SES was noted in the 0-back condition, which has low WM demands, and mainly serves as a control condition or measure of sustained attention (Miller et al. 2009). While these findings are consistent with prior work from this sample documenting greater activation associated with higher WM load (Satterthwaite et al. 2013), they contrast with another study reporting that low income adolescents recruit certain regions of the executive system, including the bilateral medial frontal gyrus and intraparietal sulcus, during low-load 0-back trials, whereas high income adolescents recruit these regions only in high-load trials (Finn et al. 2017). These divergent results suggest that future work examining the relationship between SES and WM performance should manipulate WM load over a range of task difficulties to identify load-specific effects.

The current study replicates previous work demonstrating positive associations between SES and performance on a WM task (Noble et al. 2007) and extends prior findings by identifying executive system activation as a significant mediator of this relationship. The findings from our mediation analysis are consistent with other work that identified activation of the middle, inferior, and superior frontal gyri during a WM task as mediators of individual measures of SES and WM performance (Finn et al. 2017; Rosen et al. 2018). However, our use of high dimensional mediation analysis allowed for a data driven approach that uncovered distinct multivariate patterns of brain activation that mediated the relationship between SES and WM performance. This approach facilitates the identification of neural mediators that may be contingent on other brain regions, and allows the detection of mediators encoded in distributed patterns of activation (Geuter et al. 2020). We identified two patterns of activation that mediated the relationship between SES and WM performance: the first in the medial prefrontal cortex and left and right DLPFC, and the second in the superior frontal gyrus, right middle frontal gyrus, and frontal pole. The DLPFC has previously been linked to performance on high-load WM tasks in lesion studies (Volle et al. 2008; Barbey et al. 2013), as well as in studies of transcranial magnetic stimulation during WM tasks (Brunoni and Vanderhasselt 2014). Furthermore, these results align with previous findings that DLPFC volume mediated differences in SES and executive function in white adults (Shaked et al. 2018).

The multivariate mediation analysis revealed activation patterns that were localized to the right hemisphere, particularly in the right DLPFC (in PDM2). Previous work has demonstrated laterality differences among domains of WM, such that neural function related to verbal WM is lateralized in the left hemisphere, and spatial WM is lateralized in the right hemisphere (Walter et al. 2003; Nagel et al. 2013). The fractal *n*-back task used in the current work was designed to test non-verbal WM and elicit higher levels of spatial processing (Ragland et al. 2002). As such, the right hemisphere laterality we observed may reflect individual differences related to spatial processing required by task demands. Other studies have found that children of parents with higher status jobs showed greater left-hemisphere lateralization during auditory language tasks (Raizada et al. 2008), as well as relationships between maternal education and higher levels of lateralization during a phonological processing task in the left inferior frontal gyrus (Younger et al. 2019). Findings in the current work support and expand on these studies of verbal WM, providing evidence for mirrored laterality for visual-spatial types of WM in the right hemisphere. Additionally, they suggest that the relationship between SES and brain laterality may contribute to the observed partial mediation effect.

Notably, our approach used measures of SES defined at the block group (“neighborhood”) level, which allowed us to capture facets of an individual’s circumstances beyond their immediate family or household environment, yet at a finer geographic resolution than the more commonly used ZIP code. Previous work in public health has found that neighborhood-level factors predict health outcomes over and above measures at the individual level, like chronic kidney disease (Merkin et al. 2007) or coronary heart disease (Pollack et al. 2012). The current data emphasize that neighborhood-level indicators of SES are important to the study of neurocognitive effects as well. Importantly, we found that neighborhood-level SES was associated with executive system activation over and above the effects of individual measures of adversity, such as parental education or exposure to traumatic events. These results suggest that neighborhood-level variables capture important variability not captured at the individual level. As children and adolescents spend significant time in communities outside of their immediate families, it is important that studies of brain development examine neighborhood-level variables.

This study informs the broader literature on how the neighborhood environment supports cognitive development. Studies suggest that children growing up in poorer neighborhoods have less access to higher quality community and educational resources, in terms of school funding and physical infrastructure (Macintyre et al. 1993; Evans 2004), and have parents who are less involved in their children’s education, inside and outside of the classroom setting (Benveniste et al. 2003). These relationships are consistent with proposed theories that cognitive stimulation in the home and school environment may scaffold cognitive development, including WM function (Hackman et al. 2015; Rosen et al. 2018; Rosen et al. 2019). Given these considerations, it is possible that children in low-income neighborhoods are not receiving the level of stimulation needed in order to develop higher levels of WM, which may explain the relationship presented in the current work. This hypothesis is consistent with recent work showing that protective measures, like cognitively enriched care or positive parenting, may alter the effects of early life adversity on neural measures such as brain volume (Farah et al. 2021), patterns of myelination and cortical thickness (Hong et al. 2021) or altered brain development in the temporal lobes (Whittle et al. 2017).

Certain limitations present in the current work should be noted. The data reported in this study is cross-sectional, and the sample was not obtained with the goal of studying early life environmental adversity. Furthermore, our measurement of exposure to traumatic events measures the sum of types of exposures, and cannot account for repeated exposures within a category. The sample is representative of the Greater Philadelphia area, and may not generalize to communities with differing sociodemographic characteristics. Despite strong evidence of BOLD activation during WM as a partial mediator of SES and WM performance, cross-sectional mediation relationships cannot serve as evidence of causality, and other chains of causality between neighborhood SES and cognitive function cannot be ruled out. While our measure of neighborhood SES included various important measures describing an individual’s block-level environment, we did not include measures of crime or community violence, which have been shown to vary along with other aspects of SES (Hsieh and Pugh 1993), and may have differential effects on cognitive function (Sheridan and McLaughlin 2014).

## CONCLUSION

The current study provides evidence that neighborhood-level SES is associated with executive system activation. Additionally, we identify key brain regions that mediate the relationship between neighborhood SES and cognitive performance. These results highlight the importance of neighborhood factors in shaping the executive system and underscore the importance of identifying and protecting against environmental adversity occurring at the community level that contributes to differences in executive functioning. Additionally, given the known relationship between low-SES and risk for psychopathology, (Bradley and Corwyn 2002; Peverill et al. 2021), the current reported association between SES and WM performance supports executive dysfunction as a general risk factor for diverse psychopathology (Wolf et al. 2015; Shanmugan et al. 2016). Future research may investigate targeted interventions -- including neuromodulation and community based-interventions -- that may be utilized to improve WM performance.

## Acknowledgements

This work was supported by the National Institutes of Health (T32MH014654 to B.L., F31 MH123063-01A1 to A.P., R01MH113550 to L.P., R01MH120174 to D.R.R., R01MH119185 to D.R.R., and R56AG066656 to D.RR., R01MH113565 to D.H.W., R01MH117014 to D.H.W., M.E.C., R.C.G., R.E.G., and T.M.M., R37MH125829 to T.D.S., R01MH120482 to T.D.S., R01MH112847 to T.D.S., R01MH113550 to T.D.S., and RF1MH116920 to T.D.S.); National Institute of Biomedical Imaging and Bioengineering (R01EB016061 to M.A.L., R01EB022573 to T.D.S); National Institute on Aging (R56AG066656 to D.R.R) and the Brain & Behavior Research Foundation (2020 NARSAD Young Investigator Award to L.P.)

## REFERENCES

Aneshensel CS, Sucoff CA. 1996. The Neighborhood Context of Adolescent Mental Health. J Health Soc Behav. 37(4):293–310. doi:10.2307/2137258.

Avants BB, Epstein CL, Grossman M, Gee JC. 2008. Symmetric diffeomorphic image registration with cross-correlation: evaluating automated labeling of elderly and neurodegenerative brain. Med Image Anal. 12(1):26–41. doi:10.1016/j.media.2007.06.004.

Barbey AK, Koenigs M, Grafman J. 2013. Dorsolateral prefrontal contributions to human working memory. Cortex. 49(5):1195–1205. doi:10.1016/j.cortex.2012.05.022.

Barzilay R, Calkins ME, Moore TM, Wolf DH, Satterthwaite TD, Cobb Scott J, Jones JD, Benton TD, Gur RC, Gur RE. 2019. Association between traumatic stress load, psychopathology, and cognition in the Philadelphia Neurodevelopmental Cohort. Psychol Med. 49(2):325–334. doi:10.1017/S0033291718000880.

Benveniste L, Carnoy M, Rothstein R. 2003. All Else Equal: Are Public and Private Schools Different? 1st edition. New York: Routledge.

Best JR, Miller PH, Naglieri JA. 2011. Relations between executive function and academic achievement from ages 5 to 17 in a large, representative national sample. Learn Individ Differ. 21(4):327–336. doi:10.1016/j.lindif.2011.01.007.

Boylan JM, Robert SA. 2017. Neighborhood SES is particularly important to the cardiovascular health of low SES individuals. Soc Sci Med. 188:60–68. doi:10.1016/j.socscimed.2017.07.005.

Bradley RH, Corwyn RF. 2002. Socioeconomic Status and Child Development. Annu Rev Psychol. 53(1):371–399. doi:10.1146/annurev.psych.53.100901.135233.

Brito NH, Noble KG. 2014. Socioeconomic status and structural brain development. Front Neurosci. 8:276–276. doi:10.3389/fnins.2014.00276.

Brunoni AR, Vanderhasselt M-A. 2014. Working memory improvement with non-invasive brain stimulation of the dorsolateral prefrontal cortex: A systematic review and meta-analysis. Brain Cogn. 86:1–9. doi:10.1016/j.bandc.2014.01.008.

Butler O, Yang X-F, Laube C, Kühn S, Immordino-Yang MH. 2018. Community violence exposure correlates with smaller gray matter volume and lower IQ in urban adolescents. Hum Brain Mapp. 39(5):2088–2097. doi:10.1002/hbm.23988.

Calkins ME, Merikangas KR, Moore TM, Burstein M, Behr MA, Satterthwaite TD, Ruparel K, Wolf DH, Roalf DR, Mentch FD, et al. 2015. The Philadelphia Neurodevelopmental Cohort: constructing a deep phenotyping collaborative. J Child Psychol Psychiatry. 56(12):1356–1369. doi:10.1111/jcpp.12416.

Chén OY, Crainiceanu C, Ogburn EL, Caffo BS, Wager TD, Lindquist MA. 2018. High-dimensional multivariate mediation with application to neuroimaging data. Biostatistics. 19(2):121–136. doi:10.1093/biostatistics/kxx027.

Crone EA, Wendelken C, Donohue S, van Leijenhorst L, Bunge SA. 2006. Neurocognitive development of the ability to manipulate information in working memory. Proc Natl Acad Sci. 103(24):9315. doi:10.1073/pnas.0510088103.

Demir-Lira ÖE, Voss JL, O’Neil JT, Briggs-Gowan MJ, Wakschlag LS, Booth JR. 2016. Early-life stress exposure associated with altered prefrontal resting-state fMRI connectivity in young children. Dev Cogn Neurosci. 19:107–114. doi:10.1016/j.dcn.2016.02.003.

Evans GW. 2004. The Environment of Childhood Poverty. Am Psychol. 59(2):77–92. doi:10.1037/0003-066X.59.2.77.

Farah MJ, Sternberg S, Nichols TA, Duda JT, Lohrenz T, Luo Y, Sonnier L, Ramey SL, Montague R, Ramey CT. 2021. Randomized Manipulation of Early Cognitive Experience Impacts Adult Brain Structure. J Cogn Neurosci. 33(6):1197–1209. doi:10.1162/jocn_a_01709.

Finn AS, Minas JE, Leonard JA, Mackey AP, Salvatore J, Goetz C, West MR, Gabrieli CFO, Gabrieli JDE. 2017. Functional brain organization of working memory in adolescents varies in relation to family income and academic achievement. Dev Sci. 20(5):e12450. doi:10.1111/desc.12450.

Gathercole SE, Pickering SJ, Knight C, Stegmann Z. 2004. Working memory skills and educational attainment: evidence from national curriculum assessments at 7 and 14 years of age. Appl Cogn Psychol. 18(1):1–16. doi:10.1002/acp.934.

Geuter S, Reynolds Losin EA, Roy M, Atlas LY, Schmidt L, Krishnan A, Koban L, Wager TD, Lindquist MA. 2020. Multiple Brain Networks Mediating Stimulus–Pain Relationships in Humans. Cereb Cortex. 30(7):4204–4219. doi:10.1093/cercor/bhaa048.

Gonzalez MZ, Beckes L, Chango J, Allen JP, Coan JA. 2015. Adolescent neighborhood quality predicts adult dACC response to social exclusion. Soc Cogn Affect Neurosci. 10(7):921–928. doi:10.1093/scan/nsu137.

Gur RC, Calkins ME, Satterthwaite TD, Ruparel K, Bilker WB, Moore TM, Savitt AP, Hakonarson H, Gur RE. 2014. Neurocognitive Growth Charting in Psychosis Spectrum Youths. JAMA Psychiatry. 71(4):366–374. doi:10.1001/jamapsychiatry.2013.4190.

Gur RC, Richard J, Calkins ME, Chiavacci R, Hansen JA, Bilker WB, Loughead J, Connolly JJ, Qiu H, Mentch FD, et al. 2012. Age group and sex differences in performance on a computerized neurocognitive battery in children age 8-21. Neuropsychology. 26(2):251–265. doi:10.1037/a0026712.

Gur RE, Moore TM, Rosen AFG, Barzilay R, Roalf DR, Calkins ME, Ruparel K, Scott JC, Almasy L, Satterthwaite TD, et al. 2019. Burden of Environmental Adversity Associated With Psychopathology, Maturation, and Brain Behavior Parameters in Youths. JAMA Psychiatry. 76(9):966–975. doi:10.1001/jamapsychiatry.2019.0943.

Hackman DA, Gallop R, Evans GW, Farah MJ. 2015. Socioeconomic status and executive function: developmental trajectories and mediation. Dev Sci. 18(5):686–702. doi:10.1111/desc.12246.

Hong S-J, Sisk LM, Caballero C, Mekhanik A, Roy AK, Milham MP, Gee DG. 2021. Decomposing complex links between the childhood environment and brain structure in school-aged youth. Dev Cogn Neurosci. 48:100919. doi:10.1016/j.dcn.2021.100919.

Hsieh C-C, Pugh MD. 1993. Poverty, Income Inequality, and Violent Crime: A Meta-Analysis of Recent Aggregate Data Studies. Crim Justice Rev. 18(2):182–202. doi:10.1177/073401689301800203.

Jenkinson M, Beckmann CF, Behrens TEJ, Woolrich MW, Smith SM. 2012. FSL. NeuroImage. 62(2):782–790. doi:10.1016/j.neuroimage.2011.09.015.

Leonard JA, Mackey AP, Finn AS, Gabrieli JDE. 2015. Differential effects of socioeconomic status on working and procedural memory systems. Front Hum Neurosci. 9:554. doi:10.3389/fnhum.2015.00554.

Leonard JA, Romeo RR, Park AT, Takada ME, Robinson ST, Grotzinger H, Last BS, Finn AS, Gabrieli JDE, Mackey AP. 2019. Associations between cortical thickness and reasoning differ by socioeconomic status in development. Dev Cogn Neurosci. 36:100641. doi:10.1016/j.dcn.2019.100641.

Leventhal T, Brooks-Gunn J. 2000. The neighborhoods they live in: The effects of neighborhood residence on child and adolescent outcomes. Psychol Bull. 126(2):309–337. doi:10.1037/0033-2909.126.2.309.

Luna B, Marek S, Larsen B, Tervo-Clemmens B, Chahal R. 2015. An integrative model of the maturation of cognitive control. Annu Rev Neurosci. 38:151–170. doi:10.1146/annurev-neuro-071714-034054.

Macintyre S, Maciver S, Sooman A. 1993. Area, Class and Health: Should we be Focusing on Places or People? J Soc Policy. 22(2):213–234. doi:10.1017/S0047279400019310.

Marcus D, Harwell J, Olsen T, Hodge M, Glasser M, Prior F, Jenkinson M, Laumann T, Curtiss S, Van Essen D. 2011. Informatics and Data Mining Tools and Strategies for the Human Connectome Project. Front Neuroinformatics. 0. doi:10.3389/fninf.2011.00004. [accessed 2021 Jul 19]. https://www.frontiersin.org/articles/10.3389/fninf.2011.00004/full.

Merkin SS, Diez Roux AV, Coresh J, Fried LF, Jackson SA, Powe NR. 2007. Individual and neighborhood socioeconomic status and progressive chronic kidney disease in an elderly population: The Cardiovascular Health Study. Soc Sci Med. 65(4):809–821. doi:10.1016/j.socscimed.2007.04.011.

Miller KM, Price CC, Okun MS, Montijo H, Bowers D. 2009. Is the n-back task a valid neuropsychological measure for assessing working memory? Arch Clin Neuropsychol Off J Natl Acad Neuropsychol. 24(7):711–717. doi:10.1093/arclin/acp063.

Moore TM, Martin IK, Gur OM, Jackson CT, Scott JC, Calkins ME, Ruparel K, Port AM, Nivar I, Krinsky HD, et al. 2016. Characterizing social environment’s association with neurocognition using census and crime data linked to the Philadelphia Neurodevelopmental Cohort. Psychol Med. 46(3):599–610. doi:10.1017/S0033291715002111.

Nagel BJ, Herting MM, Maxwell EC, Bruno R, Fair D. 2013. Hemispheric lateralization of verbal and spatial working memory during adolescence. Brain Cogn. 82(1):58–68. doi:10.1016/j.bandc.2013.02.007.

Noble KG, Houston SM, Brito NH, Bartsch H, Kan E, Kuperman JM, Akshoomoff N, Amaral DG, Bloss CS, Libiger O, et al. 2015. Family income, parental education and brain structure in children and adolescents. Nat Neurosci. 18(5):773–778. doi:10.1038/nn.3983.

Noble KG, McCandliss BD, Farah MJ. 2007. Socioeconomic gradients predict individual differences in neurocognitive abilities. Dev Sci. 10(4):464–480. doi:10.1111/j.1467-7687.2007.00600.x.

Peverill M, Dirks MA, Narvaja T, Herts KL, Comer JS, McLaughlin KA. 2021. Socioeconomic status and child psychopathology in the United States: A meta-analysis of population-based studies. Clin Psychol Rev. 83:101933. doi:10.1016/j.cpr.2020.101933.

Pollack CE, Slaughter ME, Griffin BA, Dubowitz T, Bird CE. 2012. Neighborhood socioeconomic status and coronary heart disease risk prediction in a nationally representative sample. Public Health. 126(10):827–835. doi:10.1016/j.puhe.2012.05.028.

Pollak SD, Wolfe BL. 2020. How developmental neuroscience can help address the problem of child poverty. Dev Psychopathol. 32(5):1640–1656. doi:10.1017/S0954579420001145.

Ragland JD, Turetsky BI, Gur RC, Gunning-Dixon F, Turner T, Schroeder L, Chan R, Gur RE. 2002. Working memory for complex figures: an fMRI comparison of letter and fractal n-back tasks. Neuropsychology. 16(3):370–379.

Raizada RDS, Richards TL, Meltzoff A, Kuhl PK. 2008. Socioeconomic status predicts hemispheric specialisation of the left inferior frontal gyrus in young children. NeuroImage. 40(3):1392–1401. doi:10.1016/j.neuroimage.2008.01.021.

Rosen ML, Amso D, McLaughlin KA. 2019. The role of the visual association cortex in scaffolding prefrontal cortex development: A novel mechanism linking socioeconomic status and executive function. Dev Cogn Neurosci. 39:100699. doi:10.1016/j.dcn.2019.100699.

Rosen ML, Sheridan MA, Sambrook KA, Meltzoff AN, McLaughlin KA. 2018. Socioeconomic disparities in academic achievement: A multi-modal investigation of neural mechanisms in children and adolescents. NeuroImage. 173:298–310. doi:10.1016/j.neuroimage.2018.02.043.

Rosenberg MD, Martinez SA, Rapuano KM, Conley MI, Cohen AO, Cornejo MD, Hagler DJ, Meredith WJ, Anderson KM, Wager TD, et al. 2020. Behavioral and Neural Signatures of Working Memory in Childhood. J Neurosci. 40(26):5090. doi:10.1523/JNEUROSCI.2841-19.2020.

Samartsidis P, Eickhoff CR, Eickhoff SB, Wager TD, Barrett LF, Atzil S, Johnson TD, Nichols TE. 2019. Bayesian log-Gaussian Cox process regression: with applications to meta-analysis of neuroimaging working memory studies. J R Stat Soc Ser C Appl Stat. 68(1):217–234. doi:10.1111/rssc.12295.

Satterthwaite TD, Connolly JJ, Ruparel K, Calkins ME, Jackson C, Elliott MA, Roalf DR, Hopson R, Prabhakaran K, Behr M, et al. 2016. The Philadelphia Neurodevelopmental Cohort: A publicly available resource for the study of normal and abnormal brain development in youth. Shar Wealth Brain Imaging Repos 2015. 124:1115–1119. doi:10.1016/j.neuroimage.2015.03.056.

Satterthwaite TD, Elliott MA, Ruparel K, Loughead J, Prabhakaran K, Calkins ME, Hopson R, Jackson C, Keefe J, Riley M, et al. 2014. Neuroimaging of the Philadelphia Neurodevelopmental Cohort. NeuroImage. 86:544–553. doi:10.1016/j.neuroimage.2013.07.064.

Satterthwaite TD, Ruparel K, Loughead J, Elliott MA, Gerraty RT, Calkins ME, Hakonarson H, Gur RC, Gur RE, Wolf DH. 2012. Being right is its own reward: load and performance related ventral striatum activation to correct responses during a working memory task in youth. NeuroImage. 61(3):723–729. doi:10.1016/j.neuroimage.2012.03.060.

Satterthwaite TD, Wolf DH, Erus G, Ruparel K, Elliott MA, Gennatas ED, Hopson R, Jackson C, Prabhakaran K, Bilker WB, et al. 2013. Functional Maturation of the Executive System during Adolescence. J Neurosci. 33(41):16249–16261. doi:10.1523/JNEUROSCI.2345-13.2013.

Shaked D, Katzel LI, Seliger SL, Gullapalli RP, Davatzikos C, Erus G, Evans MK, Zonderman AB, Waldstein SR. 2018. Dorsolateral prefrontal cortex volume as a mediator between socioeconomic status and executive function. Neuropsychology. 32(8):985–995. doi:10.1037/neu0000484.

Shamosh NA, DeYoung CG, Green AE, Reis DL, Johnson MR, Conway ARA, Engle RW, Braver TS, Gray JR. 2008. Individual Differences in Delay Discounting: Relation to Intelligence, Working Memory, and Anterior Prefrontal Cortex. Psychol Sci. 19(9):904–911. doi:10.1111/j.1467-9280.2008.02175.x.

Shanmugan S, Wolf DH, Calkins ME, Moore TM, Ruparel K, Hopson RD, Vandekar SN, Roalf DR, Elliott MA, Jackson C, et al. 2016. Common and Dissociable Mechanisms of Executive System Dysfunction Across Psychiatric Disorders in Youth. Am J Psychiatry. 173(5):517–526. doi:10.1176/appi.ajp.2015.15060725.

Sheridan MA, McLaughlin KA. 2014. Dimensions of early experience and neural development: deprivation and threat. Trends Cogn Sci. 18(11):580–585. doi:10.1016/j.tics.2014.09.001.

Sheridan MA, Peverill M, Finn AS, McLaughlin KA. 2017. Dimensions of childhood adversity have distinct associations with neural systems underlying executive functioning. Dev Psychopathol. 29(5):1777–1794. doi:10.1017/S0954579417001390.

Simmonds DJ, Hallquist MN, Luna B. 2017. Protracted development of executive and mnemonic brain systems underlying working memory in adolescence: A longitudinal fMRI study. NeuroImage. 157:695–704. doi:10.1016/j.neuroimage.2017.01.016.

Snodgrass JG, Corwin J. 1988. Pragmatics of measuring recognition memory: Applications to dementia and amnesia. J Exp Psychol Gen. 117(1):34–50. doi:10.1037/0096-3445.117.1.34.

Taylor RL, Cooper SR, Jackson JJ, Barch DM. 2020. Assessment of Neighborhood Poverty, Cognitive Function, and Prefrontal and Hippocampal Volumes in Children. JAMA Netw Open. 3(11):e2023774–e2023774. doi:10.1001/jamanetworkopen.2020.23774.

Thurstone LL. 1935. The Vectors of Mind. University of Chicago Press: Chicago (The Vectors of Mind.).

Tomlinson RC, Burt SA, Waller R, Jonides J, Miller AL, Gearhardt AN, Peltier SJ, Klump KL, Lumeng JC, Hyde LW. 2020. Neighborhood poverty predicts altered neural and behavioral response inhibition. NeuroImage. 209:116536. doi:10.1016/j.neuroimage.2020.116536.

Tooley UA, Mackey AP, Ciric R, Ruparel K, Moore TM, Gur RC, Gur RE, Satterthwaite TD, Bassett DS. 2019. Associations between Neighborhood SES and Functional Brain Network Development. Cereb Cortex. 30(1):1–19. doi:10.1093/cercor/bhz066.

Ullman H, Almeida R, Klingberg T. 2014. Structural Maturation and Brain Activity Predict Future Working Memory Capacity during Childhood Development. J Neurosci. 34(5):1592–1598. doi:10.1523/JNEUROSCI.0842-13.2014.

Vargas T, Damme KSF, Mittal VA. 2020. Neighborhood deprivation, prefrontal morphology and neurocognition in late childhood to early adolescence. NeuroImage. 220:117086. doi:10.1016/j.neuroimage.2020.117086.

Volle E, Kinkingnéhun S, Pochon J-B, Mondon K, Thiebaut de Schotten M, Seassau M, Duffau H, Samson Y, Dubois B, Levy R. 2008. The Functional Architecture of the Left Posterior and Lateral Prefrontal Cortex in Humans. Cereb Cortex. 18(10):2460–2469. doi:10.1093/cercor/bhn010.

Walter H, Bretschneider V, Grön G, Zurowski B, Wunderlich AP, Tomczak R, Spitzer M. 2003. Evidence for Quantitative Domain Dominance for Verbal and Spatial Working Memory in Frontal and Parietal Cortex. Cortex. 39(4):897–911. doi:10.1016/S0010-9452(08)70869-4.

Whittle S, Vijayakumar N, Simmons JG, Dennison M, Schwartz O, Pantelis C, Sheeber L, Byrne ML, Allen NB. 2017. Role of Positive Parenting in the Association Between Neighborhood Social Disadvantage and Brain Development Across Adolescence. JAMA Psychiatry. 74(8):824–832. doi:10.1001/jamapsychiatry.2017.1558.

Wolf DH, Satterthwaite TD, Calkins ME, Ruparel K, Elliott MA, Hopson RD, Jackson CT, Prabhakaran K, Bilker WB, Hakonarson H, et al. 2015. Functional Neuroimaging Abnormalities in Youth With Psychosis Spectrum Symptoms. JAMA Psychiatry. 72(5):456. doi:10.1001/jamapsychiatry.2014.3169.

Younger JW, Lee K-W, Demir-Lira OE, Booth JR. 2019. Brain lateralization of phonological awareness varies by maternal education. Dev Sci. 22(6):e12807. doi:10.1111/desc.12807.

